# Mutational meltdown in asexual populations doomed to extinction

**DOI:** 10.1101/448563

**Authors:** Peter Olofsson, Logan Chipkin, Ryan C. Daileda, Ricardo B. R. Azevedo

## Abstract

Asexual populations are expected to accumulate deleterious mutations through a process known as Muller’s ratchet. Lynch and colleagues proposed that the ratchet eventually results in a vicious cycle of mutation accumulation and population decline that drives populations to extinction. They called this phenomenon mutational meltdown. Here, we analyze mutational meltdown using a multi-type branching process model where, in the presence of mutation, populations are doomed to extinction. We analyse the change in size and composition of the population and the time of extinction under this model.

## 1 Introduction

“All populations are doomed to eventual extinction.” (Lynch and Gabriel, 1990)

In the absence of back mutations, an asexual individual cannot produce offspring carrying fewer deleterious mutations than itself. Indeed, it is always possible that individual offspring will accrue additional deleterious mutations. As a result, the class of individuals with the fewest deleterious mutations may, by chance, disappear irreversibly from the population, a process known as Muller’s ratchet (Muller, 1964; Felsenstein, 1974; Haigh, 1978). Successive “clicks” of the ratchet will cause the fitness of asexual populations to decline. Muller’s ratchet has been invoked to explain the evolution of sex (Muller, 1964; Felsenstein, 1974; Gordo and Campos, 2008), the extinction of small populations (Lynch and Gabriel, 1990; Lynch et al, 1993; Gabriel et al, 1993), the accelerated rate of evolution of endosymbiotic bacteria (Moran, 1996), the degeneration of Y-chromosomes (Charlesworth, 1978; Gordo and Charlesworth, 2000a), and cancer progression (McFarland et al, 2013, 2014).

Haigh (1978) argued that, in a population of constant size, the ratchet should click at a rate inversely proportional to the size of the least loaded class in the population (i.e., the number of individuals carrying the lowest number of deleterious mutations). If *k* is the lowest number of deleterious mutations present in an individual in the population, the expected size of the least loaded class at mutation-selection-drift equilibrium is

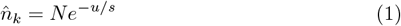

where *N* is the size of the population, *u* is the expected number of new deleterious mutations per genome per generation, and *s* is the deleterious effect of a mutation.

Haigh argued that genetic drift causes the actual value of *n*_*k*_ to deviate stochastically from 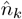. The smaller the value of 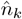, the greater the probability that *n*_*k*_ will hit zero, causing the ratchet to click. If 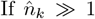, then after a click of the ratchet, the size of the new least loaded class will go to a new equilibrium, 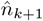, equal to 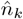 in equation (1). The value of 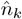 does not depend on *k* because, in a population of constant size, the evolutionary dynamics are governed by the relative fitness of individuals, not their absolute fitness (Haigh, 1978; Maia et al, 2003).

Haigh (1978) concluded that Muller’s ratchet should click faster in small populations, experiencing a high deleterious mutation rate, and mutations with milder deleterious effects (low *s*). Subsequent work has derived more accurate estimates of the rate of clicking of the ratchet using a variety of mathematical approaches (Stephan et al, 1993; Gessler, 1995; Gordo and Charlesworth, 2000a,b; Rouzine et al, 2003, 2008; Jain, 2008; Etheridge et al, 2009; Waxman and Loewe, 2010; Neher and Shraiman, 2012; Metzger and Eule, 2013). Variants of the model have explored the evolutionary consequences of variable mutational effects (Butcher, 1995; Gordo and Charlesworth, 2001; Söderberg and Berg, 2007), epistasis (Kondrashov, 1994; Butcher, 1995; Colato and Fontanari, 2001; Jain, 2008), compensatory and beneficial mutations (Wagner and Gabriel, 1990; Goyal et al, 2012; Pfaffelhuber et al, 2012; Park et al, 2018), temporal environmental heterogeneity (Wardlaw and Agrawal, 2012), population structure (Campos et al, 2006; Combadão et al, 2007; Otwinowski and Krug, 2014; Park et al, 2018; Foutel-Rodier and Etheridge, 2020), and changes in population size (Lynch and Gabriel, 1990; Melzer and Koeslag, 1991; Lynch et al, 1993; Gabriel et al, 1993; Lynch et al, 1995; Fontanari et al, 2003).

Beginning with Haigh’s foundational study, most research on Muller’s ratchet has assumed that the size of a population remains constant as deleterious mutations accumulate (e.g., Stephan et al, 1993; Gessler, 1995; Gordo and Charlesworth, 2000a; Rouzine et al, 2003; Jain, 2008; Metzger and Eule, 2013) and, therefore, soft selection (Wallace, 1975). This assumption is biologically unrealistic—if true, absolute fitness would decline relentlessly but the population would be immortal (Lynch and Gabriel, 1990; Melzer and Koeslag, 1991). Lynch and colleagues studied more realistic models where the fitness of an individual influences its fertility and populations experience density-dependent regulation (Lynch and Gabriel, 1990; Lynch et al, 1993; Gabriel et al, 1993; Lynch et al, 1995)—i.e., hard selection (Wallace, 1975). They concluded that Muller’s ratchet causes population size to decline, which accelerates the ratchet, which further reduces population size. This positive feedback results in a “mutational meltdown” that drives the population to extinction (Lynch and Gabriel, 1990; Lynch et al, 1993; Gabriel et al, 1993).

In one model, Lynch et al (1993) considered a population of asexual organisms subject to density-dependent regulation. Each individual produces *R* offspring. The number of new deleterious mutations acquired by each offspring individual is Poisson distributed with parameter *u*. The offspring then undergo viability selection such that an individual with *k* ⩾ 0 deleterious mutations has a probability of survival of

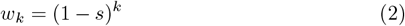

where 0 < *s* < 1 is the deleterious effect of each mutation. If the number of offspring surviving viability selection *N*′ exceeds the carrying capacity *K* then only *K* individuals survive and *N*′ − *K* individuals die, independently of their fitness; if *N*′ ⩽ *K*, all *N*′ individuals survive. Reproduction occurs after viability selection and density-dependent population regulation. Assuming that initially all individuals in the population are mutation-free and that *NR* > *K*, Muller’s ratchet proceeds in three phases in this model. First, mutations enter the population and accumulate rapidly. As the distribution of mutation numbers approaches mutation-selection-drift equilibrium (equation (1)) mutation accumulation slows down. Second, the rate of mutation accumulation settles into a steady rate. This phase proceeds as in Haigh’s model of Muller’s ratchet and lasts while 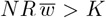, where 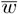 is the mean viability of the population. Third, when *w* falls below 1*/R* (i.e., when *NR w* < *K*) the population size begins to decline, triggering mutational meltdown (Lynch et al, 1993; Bull et al, 2007). Ultimately, the population goes extinct.

Lynch et al (1993) derived some analytical expressions to describe the dynamics of mutation accumulation during the first two phases and the times at which these two phases end. However, they did not present any analytical theory on the dynamics or duration of the mutational meltdown phase itself (see also Gabriel et al, 1993; Lynch et al, 1995). Recently, Lansch-Justen et al (2022) derived analytical expressions to describe the mean number of deleterious mutations, population size, and extinction time during mutational meltdown. Here we model mutational meltdown using a multi-type branching process. We analyse the change in size and composition of the population and the time of extinction under this model.

## 2 Model

### 2.1 Overview

We model a population of asexually reproducing organisms. Every generation, each individual has a probability *u* of acquiring a deleterious mutation. The mutation rate, *u*, in our model is defined differently from that in the models of Haigh (1978) and Lynch et al (1993); however, the two formulations are equivalent if *u* is small. All mutations have the same deleterious effect (*s*), do not interact epistatically, and are irreversible.

Generations are non-overlapping. We model reproduction and death using a branching process (see section 2.3). Every generation, individuals reproduce or die independently of each other and experience hard selection. The size of the population can change over time so the population can go extinct. Deleterious mutations doom the population to eventual extinction.

### 2.2 Mutational parameters

We conducted a survey of the literature looking for empirical estimates of the genomic deleterious mutation rate, *u*, and the mean deleterious effect of a mutation, *s*. The results are summarized in Figure 1. We adopted the median values of the mutational parameters (*u* = 0.02 and *s* = 0.07) as the default in this paper.

**Fig. 1.**
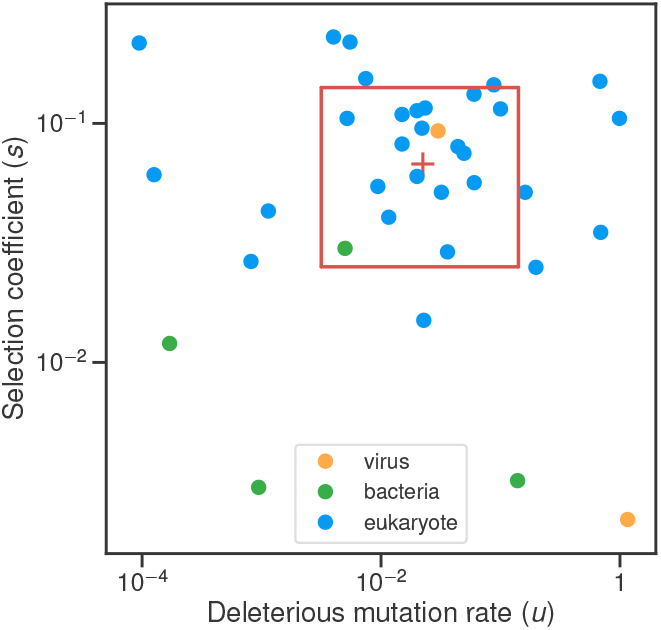
Empirical estimates of mutational parameters. We obtained 36 estimates of the genomic deleterious mutation rate (*u*) and mean deleterious effect of a mutation (*s*) from 26 studies on 16 species. When a single study included multiple estimates for a single species based on independent data sets, we used the median of those estimates. The red cross shows the median values: *u* = 0.02 and *s* = 0.07. Half of the estimates (18*/*36) are contained within the red box.

### 2.3 Branching process model

We model reproduction and death using a discrete-time, multi-type branching process (Haccou et al, 2005). We refer to individuals with *k* deleterious mutations as belonging to type *k*. Each generation, an individual of type *k* reproduces by splitting into two daughters with probability ℙ = *w*_*k*_*/*2 (Fig. 2A–C) and dies with probability ℙ = 1 − *w*_*k*_*/*2 (Fig. 2D), where *w*_*k*_ is the expected number of offspring of an individual with *k* ⩾ 0 deleterious muta-tions given by equation (2) and 0 < *s* ⩽ 1 is the deleterious effect of each mutation. Note that in our model *w*_*k*_ corresponds to fertility not viability (cf. the models of Haigh 1978 or Lynch et al 1993).

**Fig. 2.**
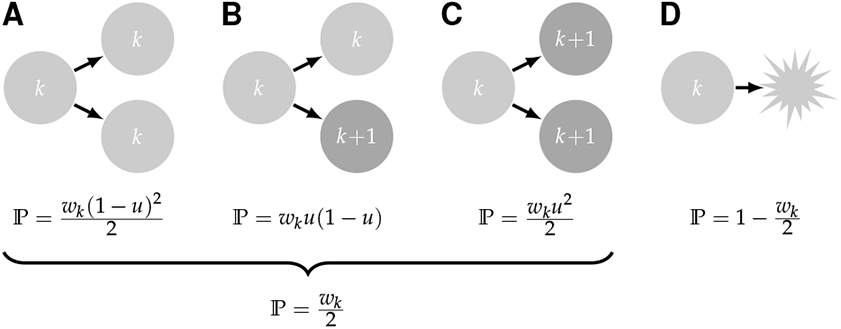
Multi-type branching process. At each time step, an individual of type *k*—i.e, with *k* deleterious mutations (light gray)—can have one of four fates (A–D) with different probabilities, ℙ; *w*_*k*_ is the absolute fitness of an individual of type *k* defined in equation (2). It can either split into two daughters (A–C) or die (D). The daughters inherit the *k* mutations from their mother. A daughter can acquire one additional mutation and become an individual of type *k* + 1 (dark gray) (B–C).

Individual offspring may independently acquire one deleterious mutation with probability *u*. The number of mutant offspring of a surviving individual of any type is, therefore, binomially distributed with parameters 2 and *u* (Fig. 2A–C). As the mean number of offspring per individual is less than 1, the process is subcritical and will go extinct with probability 1 (Haccou et al, 2005).

Our model is an example of an irreducible multi-type branching process with countably infinite type space; for an overview and examples of such processes, see Kimmel and Axelrod (2015). Pénisson et al (2013, 2017) have analyzed models similar to ours to investigate how the frequency of new alleles (neutral, deleterious, or beneficial) arising in an asexual population is influenced by later accumulation of deleterious mutations.

### 2.4 Size and composition of the population

Initially (i.e., at generation *t* = 0), a population is described by the vector

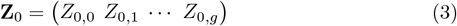

where *Z*_0,*k*_ is the number of individuals with *k* deleterious mutations and *g* is the greatest initial load (i.e., the maximum number of mutations carried by an individual; *Z*_0,*k*_ = 0 for *k* > *g*). The lowest initial load is *ℓ* (i.e., the minimum number of mutations carried by an individual; *Z*_0,*k*_ = 0 for 0 ⩽ *k* < *ℓ*).

At generation *t*, a population is described by the vector

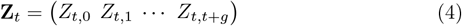

where *Z*_*t,k*_ is the number of individuals with *k* deleterious mutations. Note that *Z*_*t,k*_ = 0 for *k* > *t* + *g* because individuals can acquire no more than one mutation per generation.

In the following generation, the expected composition of the population is:

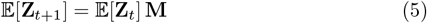

where **M** is the mean reproduction matrix with entries

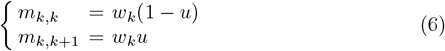

where *m*_*i,j*_ is the expected number of offspring of type *j* generated by an individual of type *i*; all other entries of **M** are 0. The expected total number of offspring of an individual of type *k* is the absolute fitness *w*_*k*_ = *m*_*k,k*_ + *m*_*k,k*+1_ (equation (2)).

Iterating (5) we get

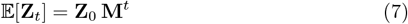

where **Z**_0_ is the (constant) initial state of the population (3). For any *t*, **M**^*t*^, the *t*-th power of **M**, is upper triangular (i.e., all its entries below the diagonal are 0). Assuming fitness function (2), we can get an explicit form for the entries of **M**^*t*^, the expected number of descendants of an individual of type *k* carrying *j additional* mutations after *t* generations:

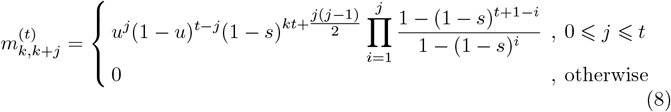

Shur (2011) provides general recursive formulas for upper diagonal matrices. Although these would apply here, in Appendix A we present an alternative proof of (8) that utilizes the particular structure of our model.

The elements of 𝔼[**Z**_*t*_] in equation (7) can be rewritten as

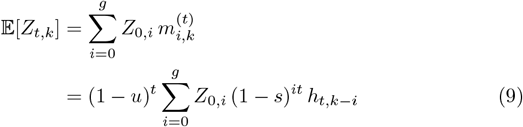

where

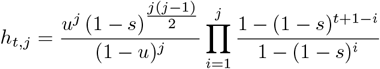

As *t* → ∞, we thus have *h*_*t,j*_ → *h*_*j*_ where

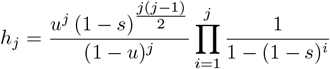

If we assume that initially there are *Z*_0,0_ mutation-free individuals in the population, equation (9) yields

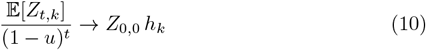

as *t* → ∞. If we further assume weak mutation (i.e., low *u*) then (1−*u*)^*t*^ ≈ *e*^−*ut*^ and we can write equation (10) as

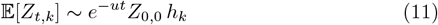

Therefore, in the long run, the numbers of individuals of all types are expected to decline exponentially at a rate equal to the mutation rate, *u*.

The total size of the population at time *t* is defined as *N*_*t*_ = *Z*_*t*,0_ + *Z*_*t*,1_ + *Z*_*t*,2_ + … + *Z*_*t,t*+*g*_; equation (9) indicates that its expected value can be computed as

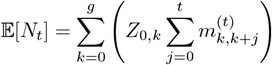

which turns out to be equivalent to

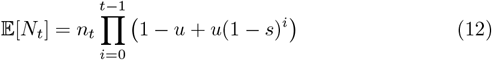

where

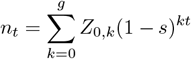

is the expected population size in the absence of mutation. For a proof of equation (12) see Appendix B.

Under weak mutation, equation (12) can be approximated asymptotically as *t* → ∞ as:

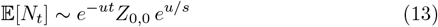

which is proved in Appendix C. Fig. 3A illustrates this approximation for default mutational parameters.

**Fig. 3.**
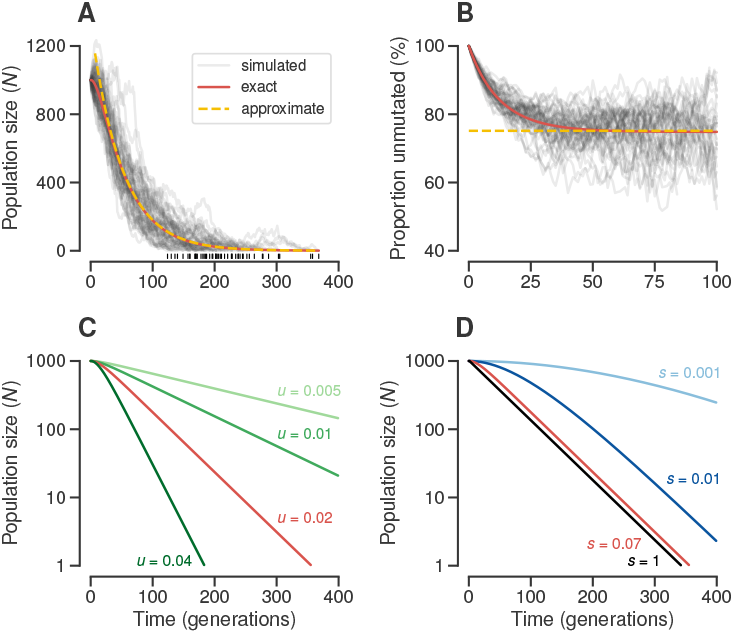
Dynamics of total population size, *N*_*t*_, during mutational meltdown. **(A)** Dynamics of *N*_*t*_ in 50 populations founded by *N*_0_ = *Z*_0,0_ = 1000 mutation-free individuals and subject to mutations with rate *u* = 0.02 and deleterious effect *s* = 0.07. Gray lines show *N*_*t*_ of individual simulated populations. The red line shows the exact expected *N*_*t*_ from equation (12). The yellow dashed line shows the long-term weak mutation approximation from equation (13). Black vertical lines above the time axis indicate extinction times. **(B)** Dynamics of the proportion of mutation-free individuals (*Z*_*t*,0_*/N*_*t*_) in the simulations shown in (A). The red line shows the ratio of the exact expected values given in equations (9) and (12). The yellow dashed line shows the long-term weak mutation approximation from equation (14). **(C)** Effect of *u* on the dynamics of *N*_*t*_ (*N*_0_ = *Z*_0,0_ = 1000, *s* = 0.07). **(D)** Effect of *s* on the dynamics of *N*_*t*_ (*N*_0_ = *Z*_0,0_ = 1000, *u* = 0.02). Note that the *N*_*t*_ axes in (C) and (D) are log-transformed. The red lines in all plots correspond to the default mutational parameters (*u* = 0.02, *s* = 0.07).

By equations (11) and (13), we thus get the ratio

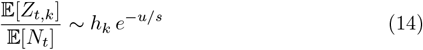

which can be viewed as a quasi-stable distribution over the classes *k* = 0, 1, …, as *t* → ∞ (“quasi” because all classes are bound to die out eventually). In particular, as *h*_0_ = 1, the mutation-free class has relative size *e*^−*u/s*^, the same proportion as predicted under the Haigh model (Haigh, 1978; Maia et al, 2003). Fig. 3B illustrates this approximation for default mutational parameters.

The computations above tacitly assume that initially the population includes some mutation-free individuals (i.e., *Z*_0,0_ > 0). However, if the least loaded type (i.e., the lowest number of mutations carried by an individual in the population) is not 0 but some number *ℓ* > 0, the ratio in equation (14) still holds with *h*_*k*_ replaced by *h*_*k*−*ℓ*_ for *k* ⩾ *ℓ*. This is easily realized by replacing (1 − *u*)^*t*^ by the product (1 − *u*)^*t*^(1 − *s*)^*ℓt*^ in the proof in Appendix C. Thus, if the ratchet clicks, the distribution will simply be displaced by one mutation (e.g., from *ℓ* to *ℓ* + 1). Haigh (1978) described a similar phenomenon in his model of Muller’s ratchet under constant population size.

### 2.5 Time to extinction

By well-known results from the theory of branching processes (Haccou et al, 2005), if the mean number of offspring per individual is strictly less than 1, the extinction time of the population has a finite expected value, 𝔼[*T*]. By equation (2), an unmutated individual has *w*_0_ = 1 offspring on average.

Equation (6), however, shows that *m*_*k,k*_ < 1 for any *k* ⩾ 0, that is, individuals of any type have on average less than one offspring of their own type, provided that mutations occur and are deleterious (*u* > 0 and *s* > 0). Therefore, the population has a finite expected extinction time.

An individual of type *k* may have offspring of types *k* or *k* + 1 according to the joint probability generating function (p.g.f.)

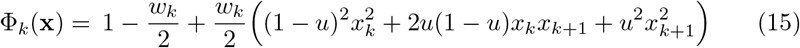

with *k* = 0, 1, … where **x** = (*x*_0_, *x*_1_, …). Note that although **x** is of infinite length, each Φ_*k*_ depends on only finitely many coordinates. Denote the joint p.g.f. of **Z**_1_ by

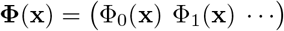

By standard branching process theory, the joint p.g.f. of **Z**_*t*_ is given by the *t*-fold composition of **Φ** with itself

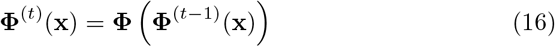

with *t* = 1, 2,, **Φ**^(0)^(**x**) = **x**, and **Φ**^(1)^(**x**) = **Φ**(**x**).

More standard theory yields that the probability that the population is extinct *by* generation *t* is

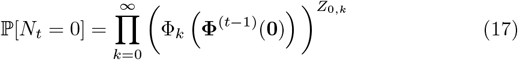

where **0** = (0, 0, 0,), and Φ_*k*_(**x**) and **Φ**^(*t*−1)^(**0**) are computed by equations (15) and (16), respectively (Mode, 1971). The probability that the population goes extinct *in* generation *t* is ℙ[*N*_*t*_ = 0] - ℙ[*N*_*t*-1_ = 0]. Fig. 4 illustrates the use of equation (17) to obtain the distribution of extinction times under default mutational parameters.

**Fig. 4.**
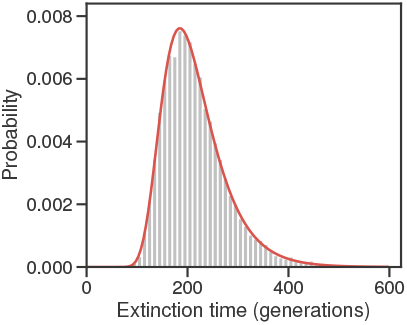
Distribution of extinction times, *T*. Histogram based on 10^4^ stochastic simulations with the same parameters as those used in Fig. 3A (*N*_0_ = *Z*_0,0_ = 1000, *u* = 0.02, *s* = 0.07). The red line shows exact probabilities computed using equation (17).

The expected extinction time *T* of the population can be computed by the formula

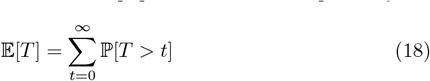

which holds for non-negative, integer-valued random variables. Moreover, as ℙ[*T* > *t*] = ℙ[*N*_*t*_ > 0], the expression becomes

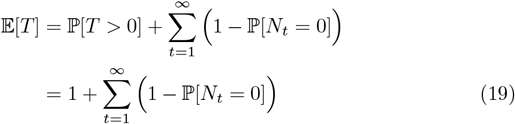

where ℙ(*N*_*t*_ = 0) is given by equation (17), and ℙ[*T* > 0] = 1 because *N*_*t*_ > 0 at time *t* = 0.

Finally, the variance of *T* is given by,

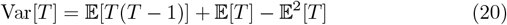

where

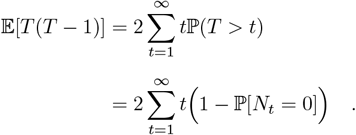

Fig. 5 uses equations (19) and (20) to compute the expected value and variance of extinction time *T* under a wide range of values of the mutational parameters. High values of the deleterious mutation rate, *u*, and selection coefficient of the mutations, *s*, both accelerate extinction (Fig. 5A). However, over the range explored *u* has a much larger effect on 𝔼[*T*] than *s*. For example, at the default selection coefficient *s* = 0.07, reducing the mutation rate by two orders of magnitude (from *u* = 0.25 to 0.0025) causes extinction times to increase 27-fold. In contrast, at the default mutation rate *u* = 0.02, reducing the selection coefficient by two orders of magnitude (from *s* = 1 to 0.01), causes extinction times to increase by only 36%. Low values of *u* and high values of *s* make extinction times more variable (Fig. 5B). Again, *u* has a stronger effect on the variability of *T* than *s*.

**Fig. 5.**
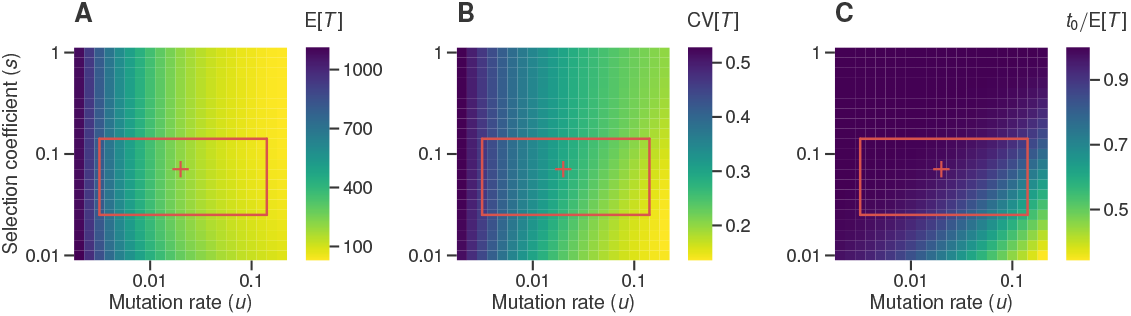
High deleterious mutation rate accelerates extinction during mutational meltdown. **(A)** Expected extinction time, 𝔼[*T*], of populations founded by *N*_0_ = *Z*_0,0_ = 1000 mutation-free individuals and subject to mutations with deleterious effect *s* and rate *u*. 𝔼[*T*] was calculated using equation (19) for 21 × 21 = 441 combinations of values of *s* and *u* evenly spaced on a log-scale spanning 2 orders of magnitude. **(B)** Coefficient of variation of extinc-tion time, 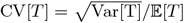 for the populations shown in (A). Var[*T*] was calculated using equation (20). **(C)** Expected extinction time of the mutation-free class, *t*_0_, as a pro-portion of 𝔼[*T*] for the populations shown in (A). *t*_0_ was calculated using equation (21). The red box and cross are described in Fig. 1.

### 2.6 The first click of the ratchet

Let *ℓ* be the least loaded class at time *t* = 0 so that initially the population includes a fixed number *Z*_0,*ℓ*_ > 0 of least loaded individuals (ancestors). Because of the irreversible nature of mutations in this model, once the least loaded class goes extinct, it can never reappear.

Let *τ*_*ℓ*_ denote the time of the first click of the ratchet during mutational meltdown (i.e., the extinction of type-*ℓ* individuals), that is

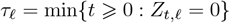

By the standard results from probability theory introduced in equations (18) and (19)

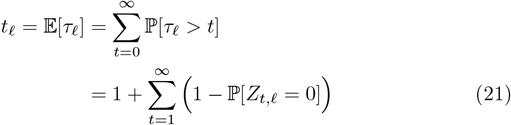

where ℙ[*τ*_*ℓ*_ > 0] = 1 because least loaded individuals are present at time *t* = 0. Fig. 5C shows that the time of first click *t*_0_ makes up a substantial proportion of the total extinction time except when *u/s* is large. For example, at the default mutational parameters, the expected time of first click is *t*_0_ = 209.2, that is, 97.3% of the total time to extinction.

The time of extinction of the entire type-*ℓ* subpopulation is the time of extinction of the *Z*_0,*ℓ*_ independent subpopulations started from the ancestors. Thus, let 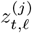 denote the number of type-*ℓ* individuals in generation *t* stemming from the *j*th ancestor and 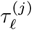 be the time of extinction of the subpopulation started from the *j*th type-*ℓ* ancestor, *j* = 1, 2, …, *Z*_0,*ℓ*_. We then have

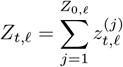

and

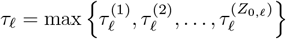

and the equivalence

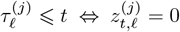

An individual of type *k* has offspring of type *k* according to the the p.g.f.

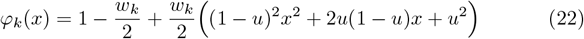

Note that *φ*_*k*_(*x*) is obtained by applying Φ_*k*_(**x**) to *x*_*k*_ in equation (15) and ignoring *x*_*k*+1_. By another standard result (Jagers, 1994), the p.g.f. of *Z*_*t,k*_ is given by the *t*-fold composition of *φ*_*k*_ with itself, denoted by 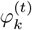.

For *t* > 0 we get the probability in equation (21)

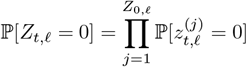

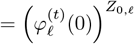

Let *X*_*ℓ*+1_ be the number of individuals of type *ℓ* + 1 present after the first click of the ratchet (i.e., the time *τ*_*ℓ*_ of extinction of type *ℓ*). *X*_*ℓ*+1_ is a random variable on *{*0, 1, 2, … *}*, noting in particular that *X*_*ℓ*+1_ may be 0, in which case both types *ℓ* and *ℓ* + 1 are extinct. The expected value of *X*_*ℓ*+1_ is given by

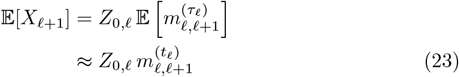

Note that equation (23) is a first-order Taylor approximation. For a proof of equation (23) see Appendix D.

### 2.7 Large least loaded class

The expected time to extinction of the least loaded class, *t*_*ℓ*_, is given by equation (21). Following Jagers et al (2007), there exists a sequence *c* (*Z*_0,*ℓ*_) → *c* as *Z*_0,*ℓ*_ → ∞ such that

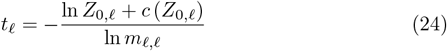

where *m*_*ℓ,ℓ*_ < 1 is the expected number of least loaded offspring per least loaded individual (equation (6)) and *Z*_0,*ℓ*_ is the initial size of the least loaded class. Note that the value of *c* depends on the parameters.

Equation (24) shows that *t*_*ℓ*_ grows logarithmically with *Z*_0,*ℓ*_ with a slope of − 1*/* ln *m*_*ℓ,ℓ*_. If mutation and selection are both weak, the slope becomes ≈ 1*/*(*u* + *ℓs*). Thus, increasing the initial size of the least loaded class delays its extinction more when mutation and selection are weak than when they are strong.

We now investigate the limiting behavior of 𝔼[*X*_*ℓ*+1_] as *Z*_0,*ℓ*_ → ∞. By equations (8), (23) and (24) we get

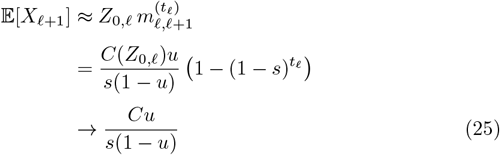

as *Z*_0,*ℓ*_ → ∞, where 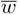 and *C* = *e*^−*c*^. For a proof, see Appendix D. Interestingly, equation (25) shows that 𝔼[*X*_*ℓ*+1_] approaches a constant as *Z*_0,*ℓ*_ increases.

If the least loaded class is mutation-free (*ℓ* = 0), the value of *t*_0_ is not affected by the effects of mutations, *s* (equations (21) and (24)), because the rate at which individuals “leave” the mutation-free class is independent of *s*. The selection coefficient does, however, affect the size of the new least loaded class, 𝔼[*X*_1_] (equations (23) and (25)), and therefore the total time to extinction.

## 3 Discussion

Most models of Muller’s ratchet have assumed that populations maintain a constant size as deleterious mutations accumulate (e.g., Haigh, 1978; Gessler, 1995; Gordo and Charlesworth, 2000a,b; Rouzine et al, 2003; Jain, 2008; Metzger and Eule, 2013). This is typically justified as resulting from density-dependent regulation of population size. However, the assumption is unrealistic because it prevents populations from ever going extinct (Lynch and Gabriel, 1990; Melzer and Koeslag, 1991). In a series of studies relaxing the assumption of constant population size, Lynch and colleagues argued that Muller’s ratchet eventually generates a positive feedback where the ratchet clicks, which causes population size to decline, which strengthens genetic drift relative to natural selection, which in turn accelerates the ratchet (Lynch and Gabriel, 1990; Lynch et al, 1993; Gabriel et al, 1993). They called this vicious cycle mutational meltdown and concluded that it drives populations to extinction.

Until recently, there was no quantitative theory of the mutational meltdown phase. We believe this theoretical neglect is unwarranted. Mutational meltdown offers a last chance for the population to be rescued by beneficial mutations or a change in the environment and avoid extinction. Thus, the dynamics and duration of the mutational meltdown phase are expected to be important determinants of the probability of evolutionary rescue (Orr and Unckless, 2008; Martin et al, 2013; Orr and Unckless, 2014; Azevedo and Olofsson, 2021).

Lansch-Justen et al (2022) derived analytical expressions to describe the mutational meltdown phase in a model similar to that originally analyzed by Lynch et al (1993). Briefly, Lansch-Justen et al began by deriving the mean number of deleterious mutations in a population assuming that natural selection is no longer effective. They then used this number to derive the expected population size through time, *N*_*t*_—analogous to our equation (12)— and then estimated time to extinction as the *t* at which *N*_*t*_ drops below 1. Lansch-Justen et al (2022) found that their approximation for extinction time matched simulation results well.

The approach described by Lansch-Justen et al (2022) does not, however, work well in our model. For example, the (exact) expected time to extinction under default mutational parameters is 𝔼[*T*] = 214.9 generations according to equation (19) (Fig. 4). The variance is Var[*T*] = 3913.6 (equation (20)). Under the same parameters, the time required for the expected population size to drop below 𝔼[*N*_*t*_] = 1 is *t* = 357 generations according to equation (12). This time overestimates the true value of 𝔼[*T*] by over two standard deviations. Furthermore, the expected population size at 𝔼[*T*] is over 17 individuals. This discrepancy is understandable because 𝔼[*N*_*t*_] takes into account populations that have already gone extinct; the probability that a population has gone extinct after *t* = 357 generations is 97% (equation 17; Fig. 4).

The extent to which real populations undergo mutational meltdown is unclear. Models of Muller’s ratchet in populations of constant size have identified three major risk factors that can drive populations into the mutational meltdown regime: long-term reductions in population size, increases in mutation rate, and intermediate deleterious effects of mutations. Next, we consider each risk factor in turn.

Population size can decline as a result of changes in the environment, such as, climate change, decreased food availability, emergence of infectious diseases, and habitat loss or fragmentation. For example, the emergence of Devil Facial Tumor Disease, a transmissible cancer, has caused the size of the Tasmanian devil population to decline by ∼77% within 5 years (Hawkins et al, 2006; Lazenby et al, 2018). As a result, the devils are under risk of extinction (McCallum et al, 2009). Our results indicate that increasing population size causes relatively small delays in extinction during mutational meltdown. *E*[*T*] is approximately proportional to the logarithm of initial population size (equation (24)). Similar results have been obtained in other stochastic models of population dynamics (Lande, 1993; Jagers et al, 2007).

Increased mutation rates can drive even very large populations into the mutational meltdown regime—a phenomenon known as lethal mutagenesis (Bull et al, 2007). Increases in mutation rate have been observed directly in experimental populations. For example, a population of *Escherichia coli* adapting to a constant environment evolved a mutator mutation after ∼25,000 generations that increased mutation rate by ∼150-fold (Barrick et al, 2009; Wielgoss et al, 2013). Evolution experiments have revealed that real populations can, indeed, experience increased extinction risk when the mutation rate is high. Zeyl et al (2001) allowed 12 populations of the yeast *Saccharomyces cerevisiae* with genetically elevated mutation rate to evolve and found that two of them went extinct within 2,900 generations. One of these populations went extinct shortly after a large decline in fitness. Bank et al (2016) subjected two populations of influenza A virus to gradually increasing concentrations of favipiravir, a drug that increases the mutation rate of the virus, and observed that both populations accumulated mutations rapidly and went extinct. The results from both of these studies are broadly consistent with the occurrence of a mutational meltdown.

The results described in the previous paragraph indicate that mutational meltdown might have clinical applications. Mutagenic agents are being explored as antiviral drugs (Loeb et al, 1999; Crotty et al, 2001; Pariente et al, 2001; Bank et al, 2016). Increased mutation rate in tumor cells has been found to correlate with improved outcomes for some cancers (Silva et al, 2000; Birkbak et al, 2011; Andor et al, 2016). Several inhibitors of key components of the DNA-repair and DNA damage-response machinery (e.g., PARP inhibitors, Lord et al, 2015), are currently being used to treat cancer, or are under preclinical or clinical development (Brown et al, 2017).

Models of Muller’s ratchet have shown that the rate at which the mean fitness of a population, 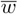, declines before entering the mutational meltdown phase is maximal at intermediate selection coefficients (Gabriel et al, 1993; Lynch et al, 1993; Gessler, 1995; Gordo and Charlesworth, 2000a,b; Lansch-Justen et al, 2022). For example, under the default mutation rate of *u* = 0.02, the decline in the 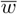 of a population of *N* = 1000 individuals is predicted to be fastest when deleterious mutations have selection coefficients of *s* = 0.0057 (decline in 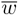 per generation was measured as *s/*Δ*t*, where Δ*t* is the time between clicks of the ratchet calculated using the method of Gordo and Charlesworth, 2000a,b). This pattern arises because when deleterious mutations have small effects selection is less efficient at purging them so the ratchet clicks faster; however, because *s* is small those clicks cause 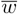 to decline slowly. Conversely, when deleterious mutations have large effects selection is efficient at purging them so the ratchet clicks slowly which also results in a slow decline of 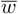. Faster declines of 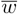 are achieved for intermediate values of *s* because selection is inefficient enough at purging deleterious mutations that the ratchet clicks relatively quickly *and* each click has a significant impact on 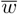. Once mutational meltdown begins, the expected extinction time decreases monotonically with *s* (Fig. 5A).

As we pointed out above, mutational meltdown offers a last chance for a population in the throes of Muller’s ratchet (Lynch and Gabriel, 1990; Lynch et al, 1993; Gabriel et al, 1993) or lethal mutagenesis (Bull et al, 2007; Matuszewski et al, 2017) to undergo evolutionary rescue (Orr and Unckless, 2008; Martin et al, 2013; Orr and Unckless, 2014; Azevedo and Olofsson, 2021). A standard model of evolutionary rescue (Orr and Unckless, 2008, 2014; Azevedo and Olofsson, 2021) considers a population that is somehow maladapted (perhaps due to sudden environmental deterioration) such that it is doomed to extinction. In the absence of beneficial mutations, this population can be modeled by a single type branching process. Models of evolutionary rescue typically ignore deleterious mutations. Interestingly, our results indicate that that simplifying assumption may be reasonable in the range of mutational parameters explored in this paper. Fig. 5A shows that for values of *u* and *s* spanning two orders of magnitude of values, the total extinction time is reasonably approximated by the case of *s* = 1, which is effectively a single type branching process.

## Acknowledgments

We thank Alex Stewart, Ata Kalirad, Herbert Levine, and Erin Kelleher for helpful discussions and Richard Neher for comments on an earlier version of the manuscript. The National Science Foundation (grants DEB-1354952 and DEB-2014566 awarded to R.B.R.A.) and National Institutes of Health (grant R01GM101352 awarded to R.B.R.A. and grant R15GM093957 awarded to P.O.) funded this work.

## Statements and Declarations

### Ethics approval

Not applicable.

### Consent to participate

Not applicable.

### Consent for publication

Not applicable.

### Availability of data and materials

Data available at https://github.com/rbazev/doomed.

### Code availability

Numerical calculations and stochastic simulations of the branching process models were done using software written in Python 3.7 and are available at https://github.com/rbazev/doomed.

### Authors’ contributions

Not applicable.

## Appendix A Proof of equation (8)

Given real numbers *b* and *x* and *n* ∈ ℕ (the nonnegative integers), let **A** denote the “almost diagonal” *n × n* matrix

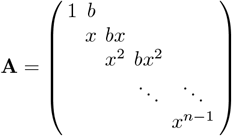

whose *i*th row is simply *x*^*i*−1^ multiplied by (0 0 … 0 1 *b* 0 0 0 …), the 1 occuring in the *i*th position. In equation (5), **M** = (1 − *u*)**A** with *x* = 1 − *s* and 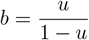.

Since **A** is upper triangular, so is its *k*th power (for *k* ∈ ℕ), with diagonal entries

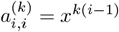

The superdiagonal entries are more complicated, but can also be expressed explicitly in terms of *x, k* and *b*.

**Lemma 1** *For n, k* ∈ ℕ, 1 ⩽ *i* ⩽ *n* − 1 *and* 1 ⩽ *j* ⩽ *n* − *i one has*

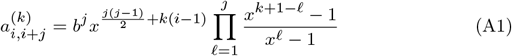

*provided that we declare x*^0^ = 1.

*Proof* We induct on *k*. When *k* = 1, for *j* = 1 and any *i* equation (A1) becomes

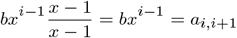

When *j* ⩾ 2, then the *ℓ* = 2 factor in the product in equation (A1) is 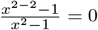, so that regardless of *i* the entire expression becomes 0, which again equals *a*_*i,i*+*j*_. We conclude that equation (A1) holds for *k* = 1.

Now assume the result is true for some *k* ⩾ 1. Since **A**^*k*+1^ = **A** *·* **A**^*k*^ and *a*_*i,ℓ*_ = 0 unless *ℓ* ∈ *{i, i* + 1*}*,

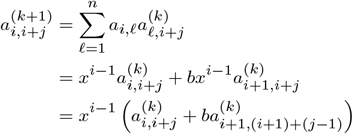

Using the inductive hypothesis^1^ we obtain

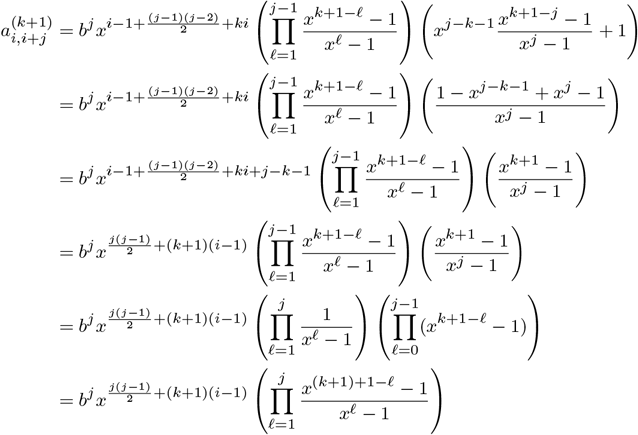

which shows that equation (A1) holds for the exponent *k* + 1. This concludes the proof.

## Appendix B Proof of equation (12)

Equation (12) provides an exact calculation of the expected population size at generation *t* given the initial state **Z**_0_:

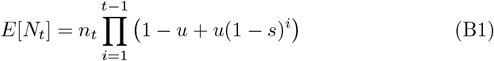

where

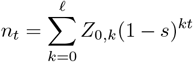

and *ℓ* is the maximum number of mutations carried by individuals present at time *t* = 0.

*Proof* Let *z*_*t,k*_ = *E*[*Z*_*t,k*_] when starting from one individual of type *k*, so that

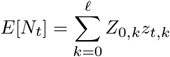

We also have

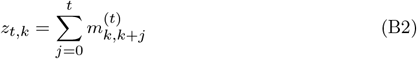

where 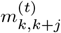 is defined in equation (8).

Equation (B1) will follow if we can prove that

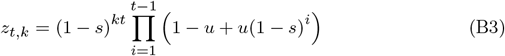

We will use induction on *t*. From equation (B2) we get 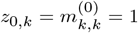 and 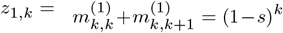, which agree with equation (B3). Now condition on the first generation. The expected number of offspring of the type-*k* ancestor is (1 − *u*)(1 − *s*)^*k*^ of type *k* and *u*(1 − *s*)^*k*^ of type *k* + 1 (equation (6)), and we get

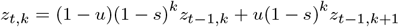

As

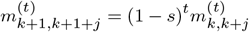

we get

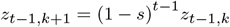

and hence the recurrence relation

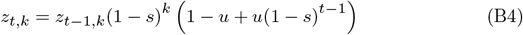

The induction hypothesis is that equation (B3) holds for *t* - 1, whence it immediately follows that it also holds for *t*.

□

## Appendix C Proof of equation (13)

Equation (13) provides the following asymptotic formula under the assumption of weak mutation (i.e., small *u*):

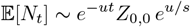

*Proof* We have

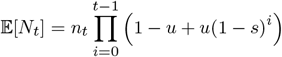

Where

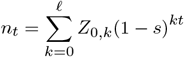

First note that because 0 < 1 − *s* < 1, we get *n*_*t*_ → *Z*_0,0_ as *t* → ∞. Next, let

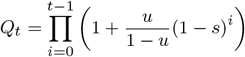

so that 𝔼[*N*_*t*_] = *n*_*t*_(1−*u*)^*t*^*Q*_*t*_. For small *u*, we use the approximation 1+*x* ≈ *e*^*x*^ to get

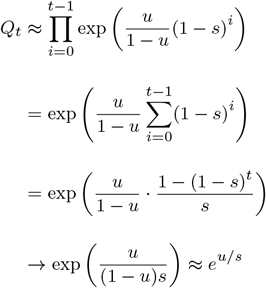

proving equation (13).

## Appendix D Proof of equation (25)

We have

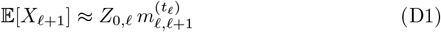

where by equation (8)

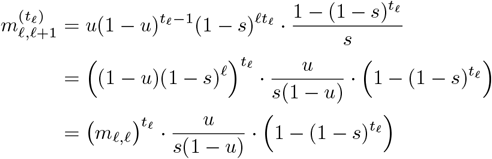

By equation (24),

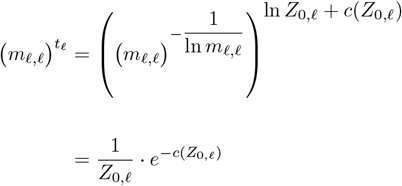

noting that, for any *a* > 0,

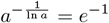

Thus, equation (D1) becomes

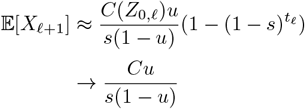

as *Z*_0,*k*_ → ∞, which is equation (25).

## Footnotes

1 Strictly speaking, the inductive hypothesis will only apply to the term 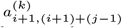 when *j* ⩾ 2. However, if we adopt the convention that any empty product is equal to one, the expression stated in the result agrees with 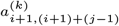 when *j* = 1 as well.

